# Defining the mechanism of galectin-3-mediated TGF-β1 activation and its role in lung fibrosis

**DOI:** 10.1101/2023.10.11.561855

**Authors:** Jessica F. Calver, Nimesh R. Parmar, Gemma Harris, Ryan M. Lithgo, Panayiota Stylianou, Fredrik R. Zetterberg, Bibek Gooptu, Alison C. Mackinnon, Robert J. Slack, Stephen B. Carr, David J. Scott, R. Gisli Jenkins, Alison E. John

**Affiliations:** School of Medicine, University of Nottingham, City Hospital Campus, Nottingham, NG5 1PB, United Kingdom; Stevenage Bioscience Catalyst, Galecto Biotech AB, Stevenage, SG1 2FX, United Kingdom; Roche Products Limited, Welwyn Garden City, Hertfordshire, AL7 1TW, United Kingdom; Research Complex at Harwell, Rutherford Appleton Laboratory, Didcot, Oxfordshire, OX11 0FA, United Kingdom; School of Biosciences, University of Nottingham, Sutton Bonington Campus, Leicestershire, LE12 5RD, United Kingdom; Membrane Protein Laboratory, Diamond Light Source, Rutherford Appleton Laboratory, Didcot, Oxfordshire, OX11 0FA, United Kingdom; Diamond Light Source, Diamond House, Rutherford Appleton Laboratories, Didcot, Oxford- shire, OX11 0FA, United Kingdom; Institute for Lung Health, NIHR Leicester Respiratory Biomedical Research Centre, Glenfield Hospital, Leicester, LE3 9QP, United Kingdom; Leicester Institute for Structural and Chemical Biology, Henry Wellcome Building, University of Leicester, Leicester, LE1 7HB, United Kingdom; Sahlgrenska Science Park, Galecto Biotech AB, Gothenburg, S-413 46, Sweden; Nine Edinburgh BioQuarter, Galecto Biotech AB, Edinburgh, EH16 4UX, United Kingdom; Department of Chemistry, University of Oxford, Oxford, Oxfordshire, OX1 3QU, United Kingdom; National Heart and Lung Institute, Imperial College London, Royal Brompton Campus, London, SW3 6LY, United Kingdom

**Keywords:** Pulmonary fibrosis, fibroblast, transforming growth factor beta (TGF-β), integrin, galectin

## Abstract

Integrin-mediated activation of the pro-fibrotic mediator transforming growth factor-β1 (TGF-β1), plays a critical role in idiopathic pulmonary fibrosis (IPF) pathogenesis. Galectin-3 is believed to contribute to the pathological wound healing seen in IPF individuals, although its mechanism of action is not precisely defined. We hypothesised that galectin-3 potentiates TGF-β1 activation and/or signaling in the lung to promote fibrogenesis. We show that galectin-3 induces TGF-β1 activation in human lung fibroblasts (HLFs) and specifically that extracellular galectin-3 promotes oleoyl-L-α-lysophosphatidic acid sodium salt (LPA)-induced integrin-mediated TGF-β1 activation. Surface plasmon resonance (SPR) analysis confirmed that galectin-3 binds to the αv integrins, αvβ1, αvβS and αvβ6 and also to the TGFβRII subunit in a glycosylation-dependent manner. This galectin-3 binding is heterogeneous and not a 1:1 binding stoichiometry. These binding interactions were blocked by small molecule inhibitors of galectin-3 which target the carbohydrate recognition domain. Binding of galectin-3 to the β1 integrin was validated *in vitro* by co-immunoprecipitation (Co-IP) in HLFs. In addition, proximity ligation assay (PLA) data indicates that galectin-3 and the β1 integrin colocalize closely (40 nm) on the cell surface of HLFs, that colocalization is increased by TGF-β1 treatment and galectin-3 inhibitors prevented this colocalization. In the absence of TGF-β1 stimulation, such colocalization was detectable only in HLFs isolated from IPF patients suggesting that the proteins are inherently more closely associated in the disease state. Taken together, this data suggests that galectin-3 promotes TGF-β1 signaling and may induce fibrogenesis by interacting directly with components of the TGF-β1 signaling cascade.

**Declaration of Interests:** RGJ reports grants or contracts from AstraZeneca, Biogen, Galecto Biotech, GlaxoSmithKline, Nordic Biosciences, RedX, Plaint, consulting fees from Bristol Myers Squibb, Chiesi, Daewoong Veracyre, Resolution Therapeutics and Pliant, honoraria from Boehringer Ingelheim, Chiesi, Roche, PatientMPower, AstraZeneca, advisory roles with Boehringer Ingelheim, Galapagos and Vicore, non-financial support from NuMedii, and is a Trustee for Action for Pulmonary Fibrosis. AEJ reports grant funding from Galecto Biotech. BG reports CASE student project partnership with Galecto Biotech, honoraria from GlaxoSmithKline and Vertex and grants and fellowships from UKRI MRC and BBSRC, Alpha-1 Foundation, BLF/A+LUK, Wellcome Trust. RJS, ACM, FRZ, JFC are Galecto employees with shares/options in the company. NRP is a Roche employee. GH, RML, PS, SBC, DJS declare no competing interests.

## Introduction

Idiopathic pulmonary fibrosis (IPF) is a chronic and progressive interstitial lung disease (ILD) (1, 2). It is the most common ILD subtype with over 32,000 patients living with the disease in the United Kingdom (UK) and more than 6,000 newly diagnosed cases annually (3). Older adults and males are primarily affected, with a median survival of 2.5-3.5 years from time of diagnosis (1, 4, 5). Repeated injury to the distal lung parenchyma of a genetically predisposed individual promotes apoptotic, necrotic and senescent pathways causing alveolar collapse and exposure of the underlying basement membrane to damage (6-9). Compromised re-epithelialisation following injury enables interstitial cells to proliferate and induces downstream signaling pathways promoting progressive lung fibrosis (10). Currently, there is no cure for IPF and only two anti-fibrotic drugs, pirfenidone and nintedanib, are licensed for IPF management (11-13).

Transforming growth factor-β1 (TGF-β1) is a potent pro-fibrotic mediator that is upregulated in IPF patients, with its protein expression concentrated to fibroblastic foci and associated with sites of active fibrosis and collagen biosynthesis (14-16). Activation of TGF-β1 via αv integrins is central to fibrogenesis, with both genetic models using ITGβ1-/- and ITGβ6-/- mice as well as anti-β6 monoclonal antibodies ameliorating bleomycin-induced lung fibrosis (17-21). Similarly in patients with IPF inhibition of αvβ6 showed promise in phase II clinical trials, although the monoclonal antibody, BG00011 (formerly STX-100) demonstrated substantial toxicity limiting its efficacy (22, 23). Therefore, alternative strategies to inhibit TGF-β1 activation and signaling are required.

Galectin-3 is a chimeric and self-associating, pro-fibrotic beta-galactoside-binding protein localised both intracellularly and extracellularly that is believed to contribute to the pathological wound healing seen in IPF individuals (24-27). Usual interstitial pneumonia (UIP) patient lung sections immunostained for galectin-3 show elevated galectin-3 expression within fibroblastic foci that is temporospatial associated with fibrosis (28). Galectin-3 levels are also higher in the serum and bronchoalveolar lavage fluid (BALF) of IPF patients compared with controls (28, 29). Following irradiation-induced lung injury *in vivo*, galectin-3 is predominately secreted by alveolar macrophages and its expression markedly increased in fibrotic lung tissue (30). Additionally, the distribution of galectin-3 is spatially related to fibrotic regions of the lung following administration of adenoviral TGF-β1 (Ad-TGF-β1) or intratracheal bleomycin *in vivo* and its mitogenic activity on cultured human lung fibroblasts (HLFs) has also been demonstrated (28, 31). Galectin-3 significantly induces fibroblast migration and increases fibroblast collagen synthesis *in vitro* further suggesting a role for galectin-3 in fibrogenesis (29). In murine models of pulmonary fibrosis, loss of galectin-3 is protective, with galectin-3-/- mice demonstrating decreased collagen staining and lower total lung collagen compared with wild type (WT) mice (28). Adding to this, the novel small-molecule inhaled galectin-3 inhibitor, GB0139 (formerly TD139), designed to modulate the fibrogenic response to tissue injury has been shown to decrease galectin-3 expression in the lung and significantly reduce bleomycin-induced fibrosis as assessed by total lung collagen content (28). In a phase 1/2a clinical trial (ID: NCT02257177) inhalation of GB0139 was shown highly suitable for dosing IPF patients and was associated with reductions in plasma biomarkers linked to IPF pathobiology including platelet-derived growth factor-beta (PDGF-BB), plasminogen activator inhibitor-1 (PAI-1), galectin-3, chemokine (C-C motif) ligand 18 (CCL18) and chitinase-3-like protein 1 (CHI3L1/ YKL-40) (32). GB0139 has since progressed into a European Union (EU) and United States (US) phase 2b clinical trial in IPF subjects ‘GALACTIC-1’ (ID: NCT03832946) with the full readout yet to the published.

Galectin-3 preferentially binds to N-glycan residues which are covalently linked to the TGF-β1 receptor during post-translation processing. Galectin-3 has been proposed to mediate cell surface retention of the TGF-β1 receptor by binding to its poly-N-acetyllactosamine residues, potentially sustaining pro-fibrotic signaling (28, 33). Similar to the TGF-β1 receptor, the literature demonstrates that galectin-3 is also capable of regulating integrin activity by interacting with their surface glycans (34-37). Therefore, we sought to define the mechanism through which galectin-3 may enhance TGF-β1 activation and contribute to the development of IPF.

## Results

### Galectin-3 induces TGF-β1 activation in fibroblasts but not epithelial cells

To investigate whether galectin-3 was able to induce TGF-β1 activation, primary HLFs and an immortalised human bronchial epithelial cell line (iHBECs) were treated with exogenous galectin-3 protein and Smad2 phosphorylation was assessed by western blotting. Although treatment with galectin-3 induced Smad2 phosphorylation in HLFs (Figure 1A), it was unable to promote phosphorylation of Smad2 in the epithelial cell line (Figure 1B). In contrast, direct stimulation with TGF-β1 was shown to induce Smad2 phosphorylation in both cell types (Figure 1A-B). Galectin-3-induced Smad2 phosphorylation in HLFs was completely inhibited by pre-treatment either with the high affinity galectin-3 inhibitor, GB0139 or with the selective inhibitor of the TGF-β type I receptor, activin receptor-like kinase (ALK5) SB-431542 (Figure 1A).

**Figure 1:**
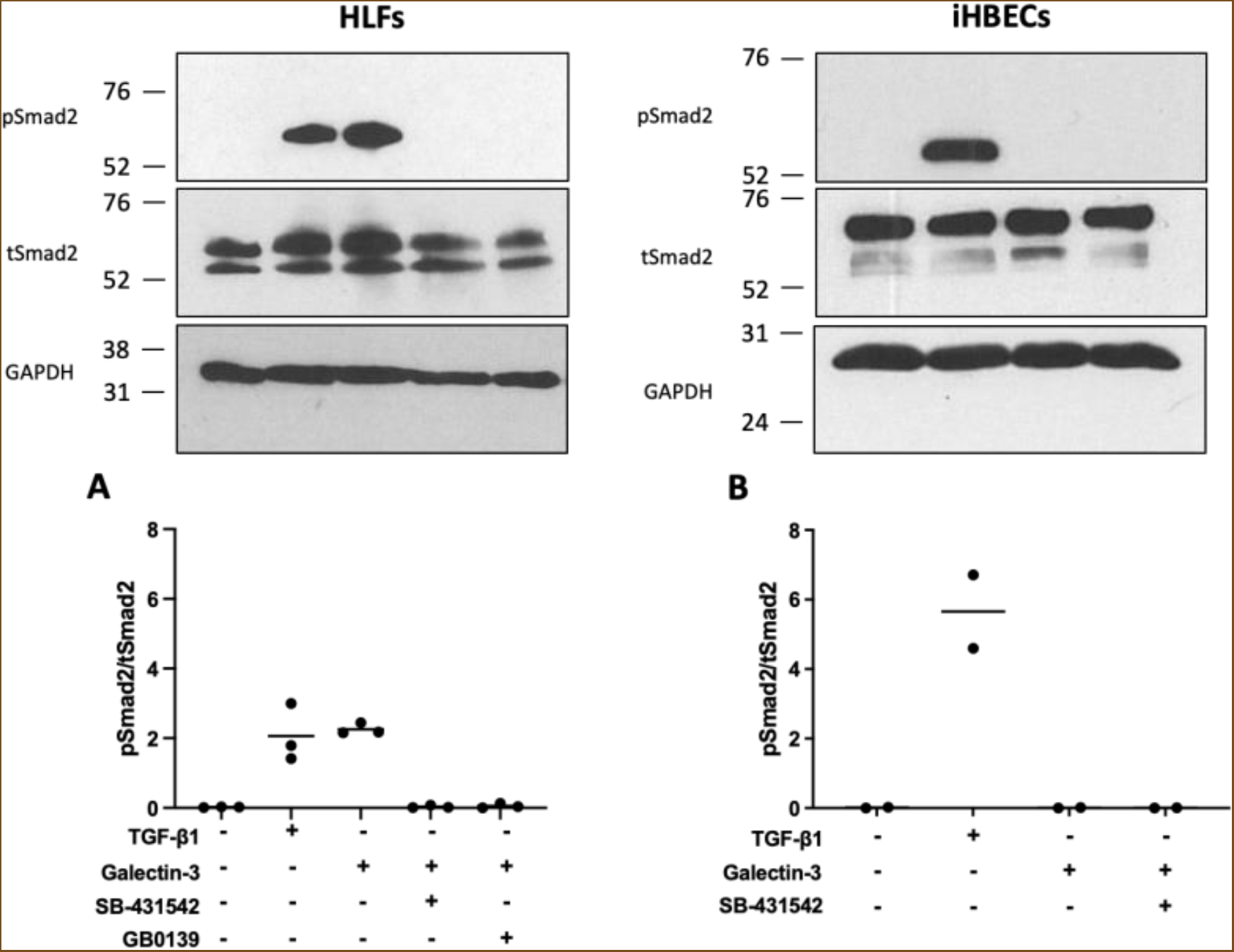
Representative western blot of pSmad2 levels in **(A)** non-IPF HLFs (N=3) and **(B)** iHBECs (N=2) pre-treated with S0 μM SB-431S42 (ALKS inhibitor) or 1 μM GB0139 (galectin-3 inhibitor) for 20 minutes prior to 2-hour treatment with 10 μg/mL galectin-3 or 2 ng/mL TGF-β1. Western blot bands were quantified using densitometry analysis and presented as a ratio of pSmad2/tSmad2.

### LPA-induced integrin-mediated TGF-β1 activation requires galectin-3

To investigate the potential involvement of galectin-3 in promoting integrin-mediated TGF-β1 activation in fibroblasts, HLFs were stimulated with oleoyl-L-α-lysophosphatidic acid sodium salt (LPA), a G-protein agonist which promotes re-organisation of the cytoskeleton and integrin activation in a variety of cell types. LPA-induced TGF-β1 activation in HLFs as measured by Smad2 phosphorylation was inhibited in a concentration-dependent manner by pre-treatment of HLFs with the β1 integrin small molecule inhibitor, NOTT199SS (Figure 2A). LPA-induced Smad2 phosphorylation in HLFs was also inhibited by pre-treatment both with the inhaled galectin-3 inhibitor, GB0139 (Figure 2B) and by the orally active galectin-3 inhibitors, GB1107 and GB1211 (Figure 2C). Pre-treatment with the cell-impermeable extracellular galectin-3 inhibitor, GB0149 (38) also inhibited phosphorylation of Smad2 in a concentration-dependent manner (Figure 2D) suggesting that galectin-3 might be mediating its effects on TGF-β1 activation extracellularly.

**Figure 2:**
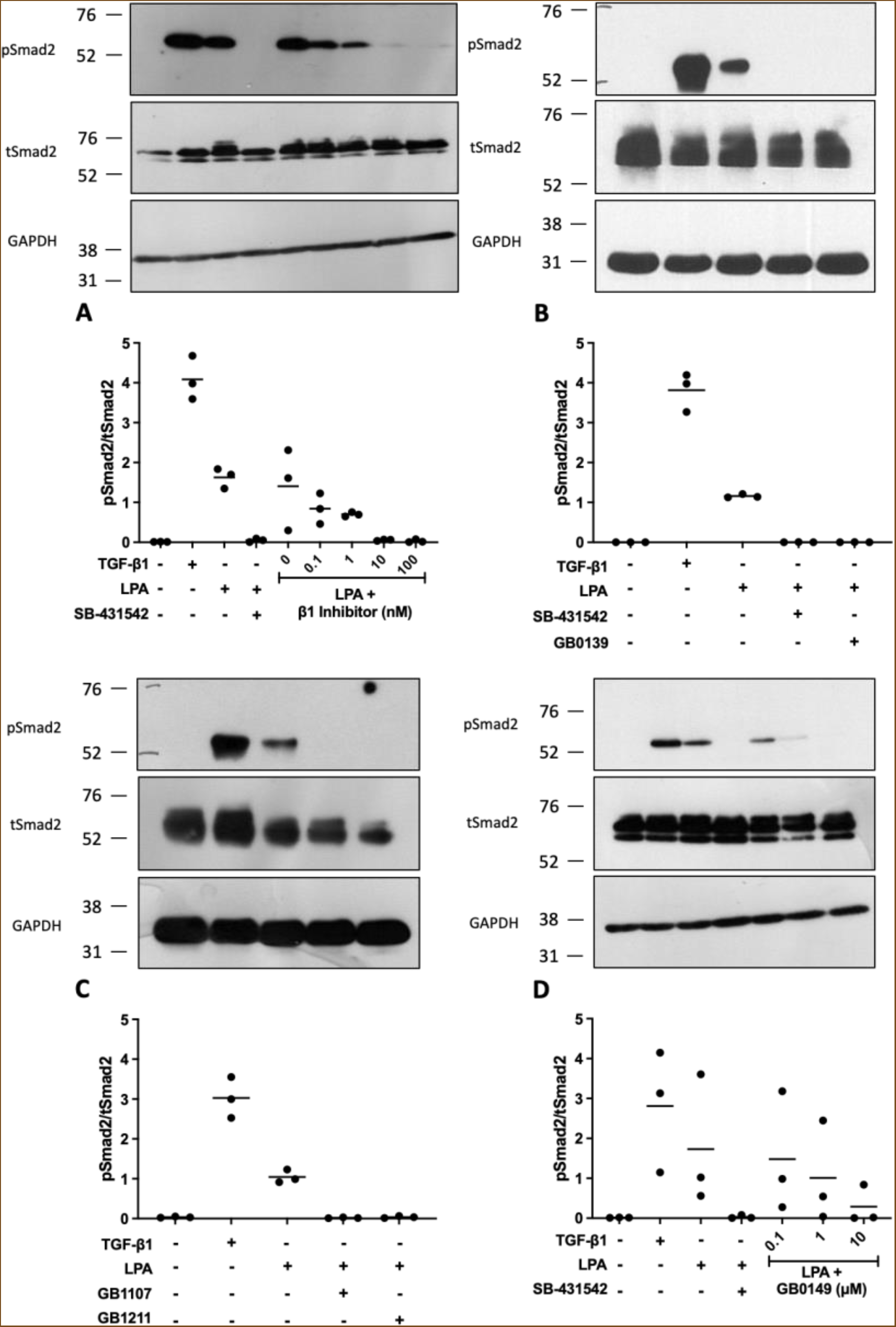
Representative western blots of pSmad2 levels in non-IPF HLFs pre-treated with **(A)** NOTT199SS β1 inhibitor (0.1-100 nM) or **(B-D)** galectin-3 inhibitors GB0139, GB1107 and GB1211 (1 μM) or GB0149 (0.1-10 μM) for 20 minutes prior to stimulation with 2 ng/mL TGF-β1 (2-hour) or 50 μM LPA (4-hour). Cells pre-treated with S0 μM SB-431542 (ALK5 inhibitor) were included as a control demonstrating maximal inhibition of pSmad2 signaling. Western blot bands were quantified using densitometry analysis and presented as a ratio of pSmad2/tSmad2.

### Galectin-3 interacts physically with integrins αvβ1, αvβ5 and αvβ6 and the TGFβRII subunit

To determine how galectin-3 promotes integrin-mediated TGF-β1 activation, protein interaction studies were performed by surface plasmon resonance (SPR). Galectin-3 bound to recombinant human integrins αvβ1 (Figure 3A), αvβS (Figure 3B) and αvβ6 (Figure 3C) in a concentration-dependent manner. Binding responses were two-fold higher with αvβ1 and αvβS compared with αvβ6 (Figure 3A-C). The highest binding responses recorded for all three integrins were considerably higher than the theoretical maximum response (approximately 135 Rmax) for analyte-ligand binding 1:1. It was not possible to saturate the immobilised integrins at the galectin-3 concentrations tested, as evidenced by the lack of a plateau. As it was not possible to yield an accurate Kd value or Bmax for these binding interactions, the minimum number of individual galectin-3 proteins involved in the binding interactions were calculated: 16, 16 and 8 for αvβ1, αvβS and αvβ6, respectively. The binding of galectin-3 to all three integrins (red lines) was seen to be glycosylation-dependent, as enzymatic removal of all N-linked and common O-linked glycans with protein deglycosylation mix II completely inhibited these binding interactions (black lines) (Figure 3A-C). Treatment with PNGase F digestion to remove N-linked glycans alone, resulted in only a partial decrease in integrin-galectin-3 binding (blue lines) (Figure 3A-C).

**Figure 3:**
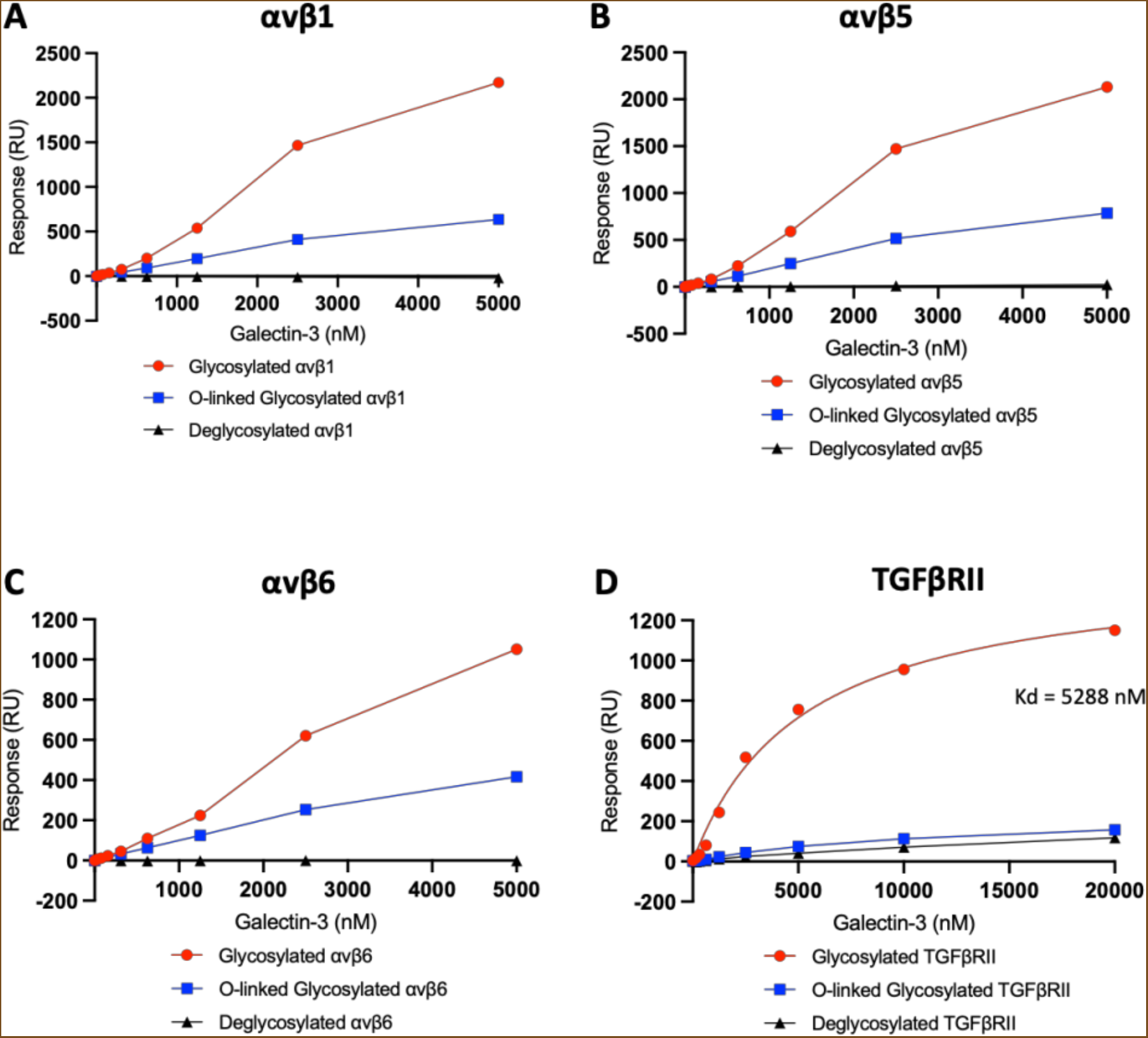
Soluble galectin-3 (sequential injections, 19.5 - 5000 nM) binding to glycosylated or deglycosylated αv integrins: **(A)** αvβ1, **(B)** αvβS and **(C)** αvβ6 immobilised on the surface of a Series S sensor chip CM5 (approximately 1000 RU). **(D)** Soluble galectin-3 (sequential injections, 156.3-20000 nM) binding to glycosylated or deglycosylated TGFβRII immobilised to a Series S sensor chip CMS (approximately 400 RU). SPR signals were measured in RU and all sensorgrams baseline-corrected. Binding response values plotted in GraphPad Prism with connecting line/curve shown.

Galectin-3 also bound to the extracellular domain of the recombinant human TGFβRII protein in a concentration-dependent manner (Figure 3D). As with the αv integrins, the highest binding response recorded was higher than the theoretical maximum response (approximately 251 Rmax) for analyte-ligand binding 1:1. As a higher analyte concentration was used for the TGFβRII experiments than for the αv integrins, saturation of the TGFβRII subunit was reached, as evidenced by the plateauing of the curve. Subsequent non-linear regression analysis yielded a Kd value of 5288 nM and a Bmax of 1473 response units (RU) and it was calculated that a minimum of 6 individual galectin-3 proteins were involved in the binding interaction. Galectin-3 binding to TGFβRII (red line) was also shown to be glycosylation-dependent with enzymatic removal of all N-linked and common O-linked glycans inhibiting the binding interaction (black line) (Figure 3D). However, a similar decrease in the level of binding was observed after the removal of only N-linked glycans (blue line) (Figure 3D) suggesting that the galectin-3-TGFβRII interaction is predominantly N-linked glycan dependent.

### Galectin-3 inhibitors prevent galectin-3 binding to αv integrins and the TGFβRII subunit

To further understand the binding of galectin-3 to integrins and the TGFβRII subunit, and the potential to disrupt this interaction with galectin-3 inhibitors, solution competition binding assays were performed. Target inhibition of the galectin-3 carbohydrate recognition domain (CRD) with GB0139 (blue line) or GB1107 (black line) inhibited the binding of galectin-3 to αv integrins: αvβ1 (Figure 4A), αvβS (Figure 4B) and αvβ6 (Figure 4C) and the TGFβRII subunit (Figure 4D) in a concentration-dependent manner. These data confirm that the interaction of galectin-3 with both the integrins and TGFβRII are CRD dependent.

**Figure 4:**
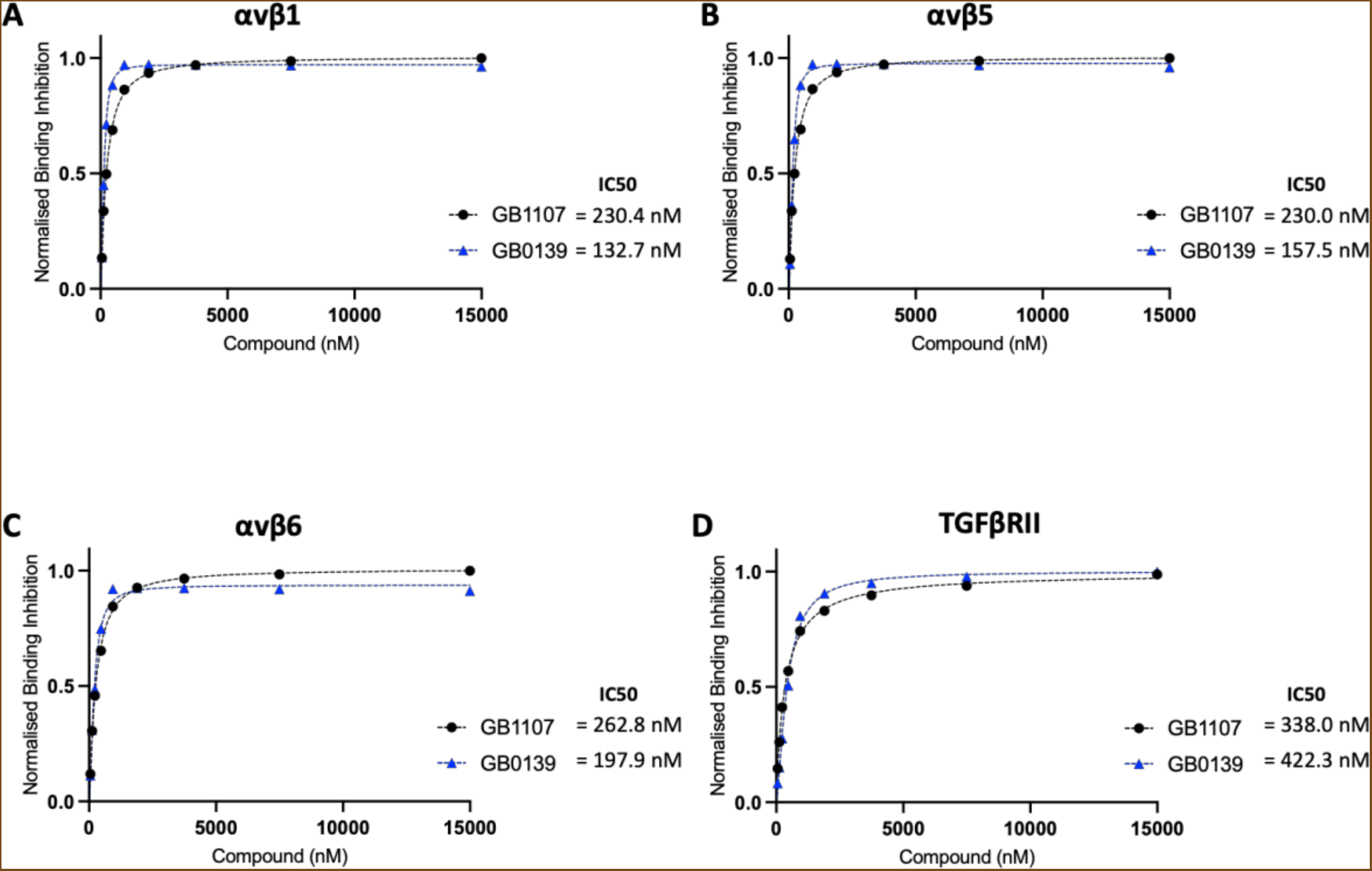
Solution competition binding assays performed with the galectin-3 inhibitor GB0139 (blue) or GB1107 (black) for αv integrins: **(A)** αvβ1, **(B)** αvβS and **(C)** αvβ6 or **(D)** TGFβRII in the presence of galectin-3 at 625 nM. Response values are normalised with respect to the highest binding response (DMSO control) and competitive inhibition graphs plotted in GraphPad Prism. IC50 values were calculated by non-linear regression analysis (binding saturation) - specific binding with hill slope.

### Galectin-3 co-immunoprecipitates with β1 integrin in HLFs

In order to confirm the key SPR binding data in a cell-based system, co-immunoprecipitation (Co-IP) experiments were performed to determine whether galectin-3 is a β1 integrin binding partner in HLFs (Figure 5). Galectin-3 (30 kDa) was successfully co-immunoprecipitated with the β1 integrin following pulldown (IP) with a β1 integrin antibody and immunoblot detection with a galectin-3 specific antibody (upper panel). No galectin-3 band was detectable in the sample immunoprecipitated with an IgG control antibody (upper panel). As expected, galectin-3 protein was detected in all input and flow through (FT) control samples (upper panel). For further confirmation of the galectin-3 and β1 integrin interaction, the Co-IP experiment was also performed in the opposite direction (lower panel). The β1 integrin (130 kDa) was successfully co-immunoprecipitated following pulldown with a galectin-3 antibody, although a low level of non-specific binding was detected in the IgG control IP (lower panel). The β1 integrin was detected in all input and FT controls (lower panel).

**Figure 5:**
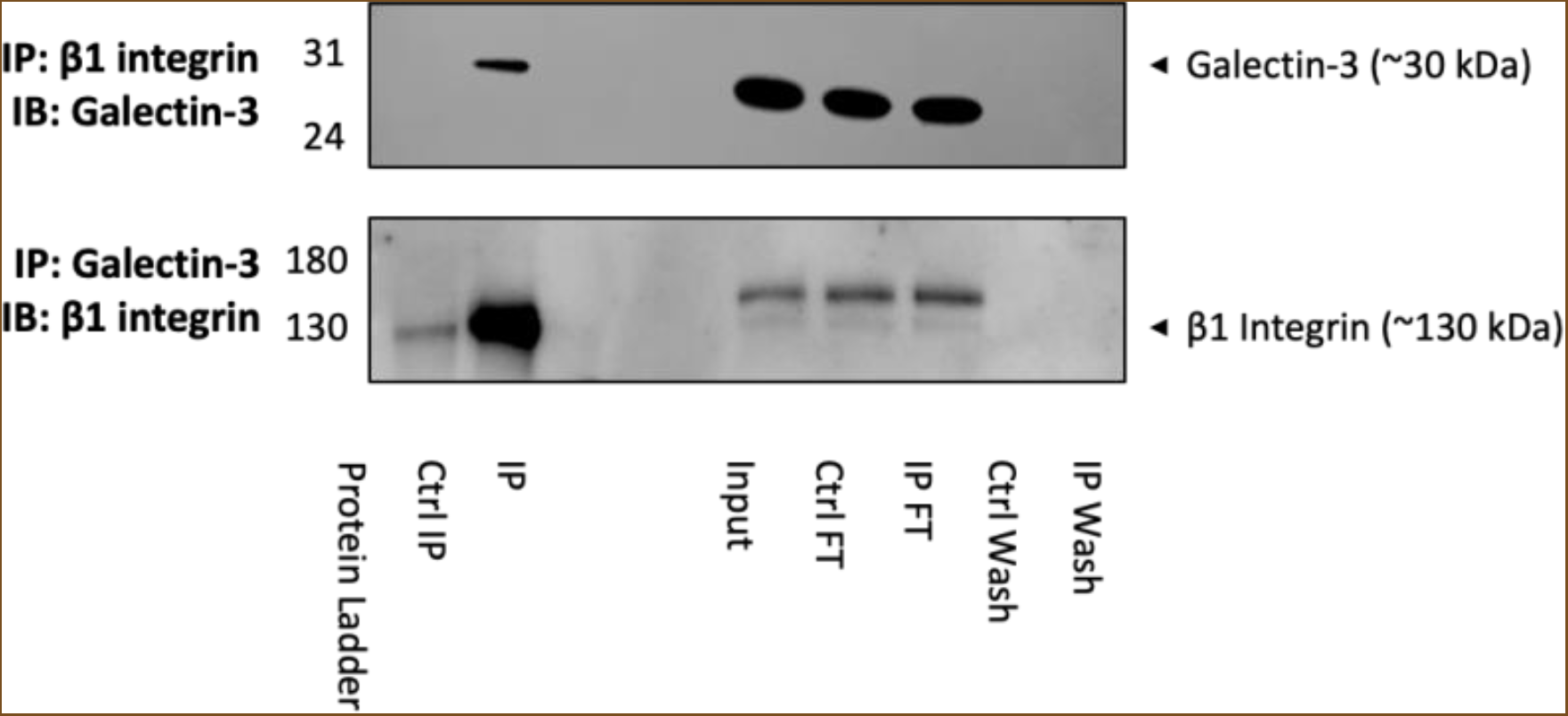
Representative western blots showing co-immunoprecipitation of galectin-3 and the β1 integrin. Whole-cell protein lysates (6S0 μg/ IP reaction) from untreated non-IPF HLFs p6 (N=3) were immunoprecipitated with an anti-β1 integrin antibody (10 μg/ IP reaction) and immunoblotted for galectin-3 (upper panel) or immunoprecipitated with an anti-galectin-3 antibody (10 μg/ IP reaction) and immunoblotted for the β1 integrin (lower panel). Co-IP input, FT and wash steps loaded as controls. Proteins separated by reducing SDS-PAGE and target protein size estimated from the marker migration pattern.

### Galectin-3 and β1 integrin are in close proximity in IPF HLFs

To further confirm that galectin-3 and the β1 integrin colocalize closely on the cell surface of HLFs, a proximity ligation assay (PLA) was performed in both non-IPF (Figure 6A) and IPF (Figure 6B) HLFs. In these experiments, the red fluorescent signal reports colocalization of galectin-3 and the β1 integrin protein within 40 nm of each other. No PLA signal was observed in untreated non-IPF HLFs (panel A1), stimulation of the non-IPF HLFs with TGF-β1 for 24 hours resulted in the appearance of a low level red fluorescence signal (panel A2). In HLFs isolated from fibrotic lung tissue higher levels of PLA signal were readily apparent. This was observed in untreated IPF HLFs (panel B1), and stimulation with TGF-β1 further augmented it (panel B2). To confirm the previous SPR and Co-IP data suggesting that compounds which target the galectin-3 CRD could prevent the interaction of galectin-3 with integrins, PLA studies were also performed in the presence of GB0139 (0.1-10 μM). Pre-treatment of the unstimulated and TGF-β1 stimulated IPF HLFs with GB0139 reduced the level of galectin-3 and β1 integrin colocalization in a concentration-dependent manner (panels B1, B3, B5, B7 for unstimulated cells, panels B2, B4, B6, B8 for cells stimulated with TGF-β1). There was abrogation of PLA signal at 1 μM and 10 μM compared to the vehicle control in both stimulated and unstimulated cells. This was also apparent in the unstimulated HLFs at 0.1 μM, but in the stimulated cells it was not clearly appreciable at this inhibitor concentration, relative to the high expression levels in the vehicle control.

**Figure 6:**
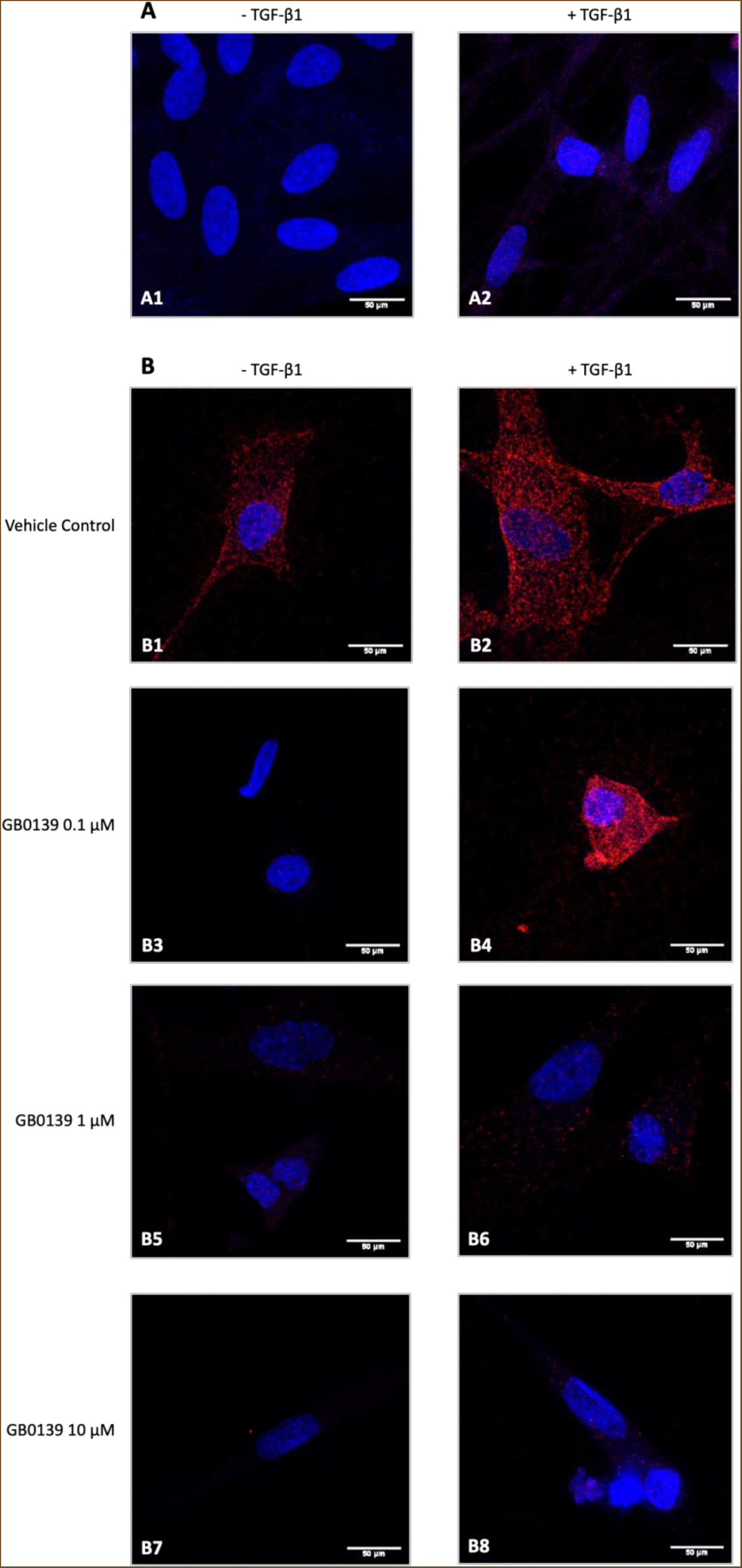
Representative confocal microscopy images (63x magnification) showing PLA of galectin-3 and the β1 integrin in **(A)** non-IPF HLFs p3-4 (N=3) or **(B)** IPF HLFs p3 (N=4) in the absence or presence of TGF-β1 stimulation (2 ng/mL TGF-β1 for 24 hours). Cells probed with a mouse anti-β1 integrin primary antibody (S μg/mL) and a rabbit anti-galectin-3 primary antibody (S μg/mL) followed by anti-rabbit PLUS and anti-mouse MINUS probes. Colocalization of galectin-3 and the β1 integrin 40 nm indicated by red fluorescence with DAPI counterstaining (blue).

## Discussion

We show that galectin-3 can activate TGF-β1 and that inhibiting galectin-3 can prevent agonist-induced integrin-mediated TGF-β1 activation in fibroblasts. These data demonstrate that galectin-3 binds integrins and the TGF-β1 receptor in a glycosylation-dependent manner. Galectin-3 binding responses were higher for αvβ1 and αvβS compared to αvβ6, which may partially explain the lack of galectin-3-mediated TGF-β1 activation in iHBECs despite high levels of αvβ6 integrin expression (39). Galectin-3 inhibition prevents binding of galectin-3 to integrins which in turn inhibits TGF-β1 activation. Given the requirement for the close physical association of the integrin, the latent TGF-β1 complex and TGF-β1 receptor for agonist-induced integrin-mediated TGF-β1 activation (40), we hypothesise that galectin-3 promotes integrin-mediated TGF-β1 activation in lung fibroblasts by facilitating clustering of the β1 integrin and TGF-β1 receptor on their respective cell surfaces (Figure 7).

**Figure 7:**
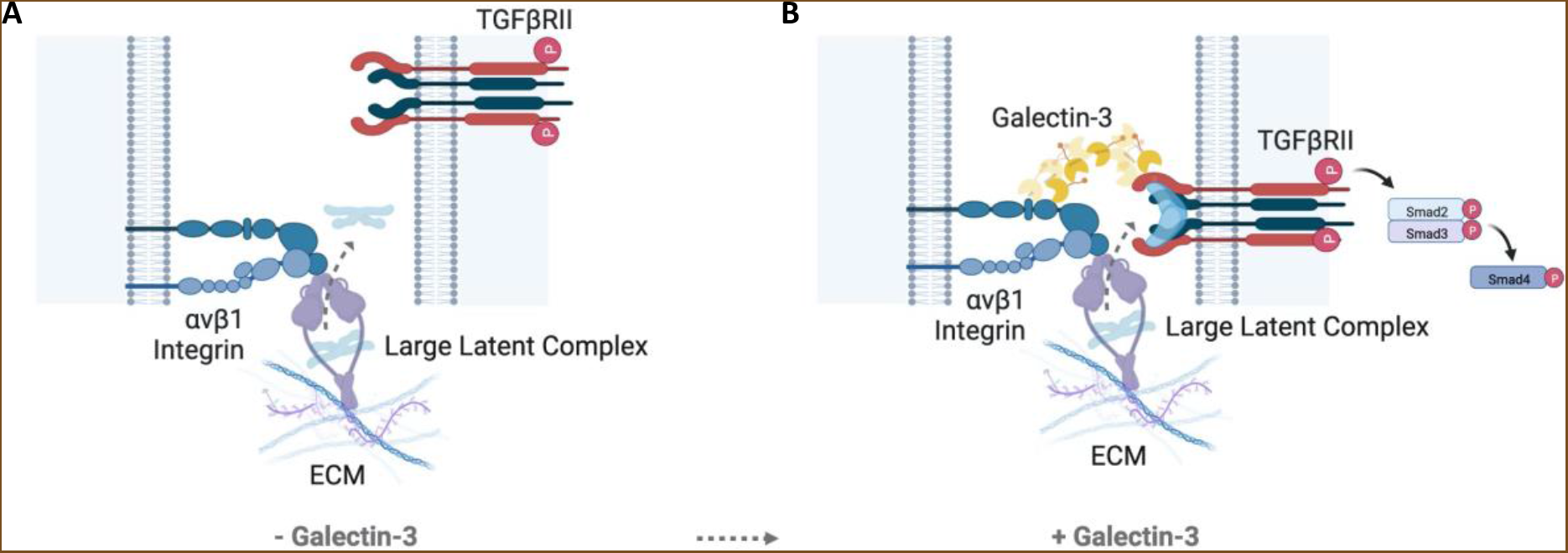
**(A)** Downstream signaling of TGF-β1 following its integrin-mediated activation requires the integrin and TGF-β1 receptor to be in close proximity on the cell surface. **(B)** The galectin-3 carbohydrate binding domain binds to the glycosylation sites on αv integrins and the TGF-β1 receptor forming a galectin lattice at the cell surface which facilitates receptor clustering. This scaffold ensures that TGF-β1 can act on its receptor and potentiates TGF-β1 signaling. GB0139 binds to the galectin-3 carbohydrate recognition domain and blocks these protein-glycan interactions.

Traction-dependent TGF-β1 activation is mediated by the integrins αvβ1, αvβ3, αvβS and αvβ6 which directly bind to the latency-associated peptide (LAP) portion of the latent TGF-β1 complex (19, 21, 41-43). The αvβ1 integrin is suggested to be the principal integrin responsible for activation of latent TGF-β1 in fibroblasts (44, 45). The phospholipid LPA has been shown to bind its G protein-coupled receptor LPA_2_ to induce αvβ6-mediated TGF-β1 activation in epithelial cells via both Ras homolog family member A (RhoA) and Rho-associated protein kinase (ROCK) (46). Although LPA has been shown to mediate primary lung fibroblast chemotaxis and proliferation via LPA_1_ following lung injury, the potential contribution of LPA signaling to integrin-mediated TGF-β1 activation in fibroblasts is unexplored (47, 48). Here we show evidence of LPA-induced αvβ1-mediated TGF-β1 activation in primary lung fibroblasts and that pre-treatment with a β1 inhibitor reduced LPA-induced Smad2 phosphorylation in a concentration-dependent manner. Subsequently, the potential significance of galectin-3 binding to αvβ1 was explored in the context of LPA-induced integrin-mediated TGF-β1 activation *in vitro*. LPA stimulation of HLFs following pre-treatment with cell-permeable galectin-3 inhibitors (GB1107 and GB1211) reduced phosphorylated Smad2, reflective of TGF-β1 activation levels. As with the β1 integrin, these findings show endogenous galectin-3 as being essential for LPA-induced TGF-β1 activation and more specifically, it is the extracellular galectin-3 that is responsible for this mechanism as the cell-impermeable galectin-3 inhibitor (GB0149) (38) also reduced Smad2 phosphorylation in a concentration-dependent manner. This role of extracellular galectin-3 was confirmed following a short incubation (20 minutes) with GB0139 which completely inhibited TGF-β1 activation as denoted by Smad2 phosphorylation. During this short time frame, target engagement of intracellular galectin-3 could not occur due to poor cell permeability (38), therefore highlighting that the effects on TGF-β1 activation are mediated by extracellular galectin-3.

Galectin-integrin binding by SPR was glycan-dependent thus supporting the view that extracellular galectin interactions are commonly carbohydrate-dependent (49-51). Enzymatic removal of all integrin N-linked and common O-linked glycans inhibited the galectin-integrin binding interaction whilst only a partial reduction in binding was observed following the removal of N-linked glycans alone. From these experiments it was not possible to determine whether the partial binding observed following PNGase F treatment was due to O-linked glycan interactions or residual N-linked glycans. Although the efficiency of PNGase F treatment was visually confirmed by a shift in protein mobility by sodium dodecyl-sulfate polyacrylamide gel electrophoresis (SDS-PAGE), this does not verify 100% N-glycan removal and therefore, some N-glycans may potentially remain on the integrin surface for galectin binding. Alternatively, the O-linked glycans may partially compensate for the higher affinity N-linked glycan interactions when they are no longer available or there is cooperative binding between the glycan subtypes. These findings are in agreement with the published literature whereby the carbohydrate dependency of galectin-integrin binding has been demonstrated using a number of techniques other than SPR (35, 37).

The galectin-3 binding responses recorded across all three integrins were considerably higher than the theoretical maximum response (approximately 135 Rmax) for analyte-ligand binding 1:1, this demonstrates that multiple individual galectins are involved in the binding interaction. As SPR measures a change in mass, it is not possible to accurately conclude if the total estimated number of galectin-3 protein is directly interacting with the immobilised ligand or if this number is a result of galectin-3 oligomerisation. As galectin-3 can oligomerise through either the N-terminal or C-terminal domain this may have resulted in multivalency of carbohydrate-binding activity where multivalent galectin-3 binds to one or more cell surface glycans causing high avidity binding (24, 25, 52-54). Alternatively, each galectin-3 protein could be binding to a single binding site on the integrin as a number of glycosylation sites have been reported or predicated from the NXS/T consensus sequence on the αv (13 sites), β1 (12 sites), βS (8 sites) and β6 subunits (9 sites) (55, 56). There is currently no consensus sequence motif to enable the prediction and identification of O-glycans. However, it is less likely that each galectin protein is binding to a single glycosylation site given the estimated number of proteins and the size of galectin-3 when compared to the integrin receptor extracellular domain. It is therefore most probable that galectin-3 binds to one or more binding sites on the integrin causing subsequent self-association and lattice formation. Supporting this, galectin-3 has previously demonstrated positive cooperativity upon binding by SPR (57).

Galectin-3 also bound to the TGFβRII extracellular domain in a glycan-dependent manner. However, in contrast with integrin-galectin-3 interactions, PNGase F treatment of TGFβRII prevented TGFβRII-galectin-3 binding to a similar level observed with total glycan-removal. This suggests that the N-linked oligosaccharides on the extracellular domain of TGFβRII predominantly mediate the galectin-3 binding interaction. This is consistent with a previous study which demonstrated alpha-1,6-mannosylglycoprotein 6-beta-N-acetylglucosaminyltransferase (Mgat5)-dependent binding of galectin-3 to the TGFβRII subunit by co-immunoprecipitation (33). In this study the half-life of the TGF-β1 receptor was reduced in Mgat5-/- cells and subsequently cytokine signaling as evidenced by decreased Smad2/3 nuclear translocation in responses to TGF-β1 (33). This suggests that N-linked glycans and their subsequent interactions are necessary for TGF-β1 canonical signaling following TGF-β1 ligand binding. Although it has been confirmed that TGFβRII has two N-linked glycan sites and a third predicted by sequence analysis (58), as with the integrins, it is not possible to conclude from this data whether galectin-3 binds to all three sites or oligomerises upon binding. These experiments do however confirm that protein-glycan binding events occur at the galectin-3 C-terminus as small molecule inhibitors of galectin-3 (GB0139 and GB1107), that act via the galectin CRD, prevented galectin-3 binding to all 3 integrins and TGFβRII in a concentration-dependent manner.

Endogenous galectin-3 bound to the β1 integrin by co-immunoprecipitation in primary lung fibroblasts, irrespective of order of IP and immunoblotting, therefore validating the positive SPR data generated with recombinant proteins. Close (40 nm) colocalization of galectin-3 and the β1 integrin was subsequently assessed by PLA in human lung fibroblasts. In the absence of TGF-β1 stimulation, such colocalization was detectable by this method in IPF HLFs but not non-IPF, suggesting that the proteins are inherently more closely associated in the disease state. These observations are consistent with non-peer reviewed results showing that colocalization of galectin-3 and the β1 integrin within 40 nm but was not detected by this method in untreated lung *ex vivo* human lung tissue (59), but was detectable in TGF-β1 treated lung tissue samples created using a validated model of lung fibrogenesis (60). Stimulation with TGF-β1 increased the signal in both cell types, more markedly in the IPF tissue-derived cells, and could be entirely abrogated by antagonism of galectin-3 glycan binding using the small molecule inhibitor GB0139. Together these findings support a mechanistic link between TGF-β1 signaling, enhanced complexation of galectin-3 with the β1 integrin via CRD:glycan interactions and fibrogenesis in disease. These data are also encouraging given that the concentration of compound used here covers the concentration range used in the phase 1/2a study and is therefore translatable to the clinic (ID: NCT02257177) (32).

Collectively, this work shows that CRD-dependent galectin-3 binding interactions are implicated in TGF-β1 signaling in human lung fibroblasts. We have performed *ex vivo* validation studies of biophysical data which confirm that galectin-3 interacts with the β1 integrin in HLFs. We have shown extracellular galectin-3 to be involved in LPA-induced integrin-mediated TGF-β1 activation and alluded to a role for galectin-3 in TGF-β1 signaling at the receptor level. We hypothesise that upon binding, galectin-3 self-associates to form a lattice between the αvβ1 integrin and TGF-β1 receptor on adjacent cells to facilitate receptor clustering. Subsequently, when TGF-β1 is activated via traction-dependent mechanisms, it is in close enough proximity to the TGF-β1 receptor to bind and signal, causing fibrogenesis. Understanding the precise role of galectin-3 in IPF pathogenesis may be critical for the continued development of more effective and selective treatments for IPF patients.

### Experimental Procedures

#### Culture of primary cells

IPF and non-IPF HLFs were obtained from explanted human lung tissue post-lung biopsy following informed, written consent and ethical review (Ethical approval numbers: 08/H0407/1, 20/SC/0142, 10/H0402/12). Briefly, the tissue was washed with PBS and then cut into 1 mm^2^ sections. Two sections were placed per well of a 6-well plate and left to adhere to the plastic for 5-10 minutes before adding culture media. Cells were cultured in Dulbecco’s Modified Eagle Medium (DMEM) high glucose (Sigma-Aldrich) supplemented with 10% foetal bovine serum (FBS) (Gibco), 4 mM L-glutamine (Sigma-Aldrich), 100 U/mL penicillin and 100 μg/mL streptomycin (Sigma-Aldrich). Cells were grown in a humidified incubator set to 37°C with 5% CO_2_.

#### Culture of immortalised cell lines

iHBECs (gifted) were cultured in keratinocyte serum-free media with L-glutamine (Gibco) supplemented with 0.2 ng/mL human recombinant epidermal growth factor (Gibco), 2S μg/mL bovine pituitary extract (Gibco), 2S μg/mL G418 disulphate (VWR) and 2S0 ng/mL puromycin dihydrochloride (Sigma-Aldrich). Cells were grown in a humidified incubator set to 37°C with 5% CO_2_.

#### Treatment of cells

Cells were growth arrested in serum-free media for 24 hours prior to stimulation in the presence or absence of inhibitors. After serum starvation, cells were stimulated with either 10 μg/mL recombinant human galectin-3 protein (R&D Systems) or 2 ng/mL recombinant human TGF-β1 protein (R&D Systems) for 2 hours or S0 μM LPA (Sigma-Aldrich) for 4 hours.

When used, inhibitors were applied in serum-free media for 20 minutes prior to stimulation. All of the small molecule glycomimetics utilised in this study, GB0139 (formerly TD139), GB1107, GB1211 or GB0149 were synthesised by the Galecto Biotech AB Medicinal Chemistry (Gothenburg, Sweden) Department (Table S-1) and used at a concentration range of 0.1-10 μM. The ALKS/ TGFβRI inhibitor SB-525334 (Sigma-Aldrich) (61, 62) was used at a concentration of S0 μM. The β1 integrin inhibitor NOTT199SS (gifted) was used at a concentration range of 0.1-100 nM. All inhibitors were dissolved in 100% dimethyl sulfoxide (DMSO), and all cells, including untreated controls were exposed to a DMSO concentration equivalent to that used in the highest inhibitor concentration in each experiment.

#### Western blot

Protein expression levels of phospho-Smad2 (pSmad2), total Smad2 (tSmad2) and glyceraldehyde 3-phosphate dehydrogenase (GAPDH) were determined by western blot as previously described (63). Briefly, cells were lysed in protein lysis buffer and protein concentrations determined by Pierce bicinchoninic acid (BCA) assay (Thermo Scientific). Protein samples (15-30 μg/lane) were subjected to SDS-PAGE on a 10% SDS-PAGE gel and electroblotted to a polyvinylidene fluoride (PVDF) membrane (Bio-Rad). After blocking for 1 hour; 5% non-fat milk in Tris-buffered saline plus 0.1% Tween 20 (TBST), the membrane was incubated overnight at 4°C with primary antibody: rabbit anti-pSmad2 mAb (Cell Signaling Technology, Clone No. 138D4; 1:1000), rabbit anti-Smad2/3 pAb (Cell Signaling Technology; 1:1000) or rabbit anti-GAPDH mAb (Abcam, Clone No. EPR16884; 1:10,000). After washes (TBST) the membrane was incubated for 1 hour at room temperature with a horseradish peroxidase (HRP)-conjugated secondary antibody: goat anti-rabbit Immunoglobulins/HRP pAb (Dako; 1:3000). All primary and secondary antibodies used were diluted in blocking buffer. The membrane was then incubated with enhanced chemiluminescence (ECL) western blot detection reagent and visualised by exposing to Hyperfilm-ECL (Thermo Fisher). Densitometry was performed using ImageJ and data plotted in GraphPad Prism as a band density ratio of pSmad2/tSmad2 to determine relative pSmad2 expression.

#### Galectin-3 expression and purification for surface plasmon resonance

The human galectin-3 gene sequence was obtained from UniProt (P17931) and delivered as plasmid DNA in a pTwist amp high copy vector (pUC origin of replication derived from pMB1 plasmid, ampicillin resistance; Twist Bioscience, USA). Galectin-3 was cloned into a pOPINF expression plasmid using infusion cloning (Addgene plasmid #26042) (64), primers (Sigma-Aldrich) used were:

5’-3’ aagttctgtttcagggcccgATGGCGGATAATTTTAGCTTACATGAC

3’-S’ atggtctagaaagctttaGATCATTGTATAGCTGGCCGAAGTG

Recombination products were used directly for the transformation of competent *E. coli* TOP10 cells (IBA Lifesciences) with carbenicillin selection (50 μg/mL). Plasmid DNA was purified using the QIAprep Spin Miniprep Kit (Qiagen) and sequence verified (Source Bioscience).

Galectin-3 was expressed in Lemo21 (DE3) cells (Thermo Fisher) in terrific broth (TB) media (carbenicillin S0 μg/mL, chloramphenicol 3S μg/mL), inoculated cultures were left to grow (37°C, 210rpm) until an OD_600nm_ of 1.4, cells were induced with 1 mM isopropylthio-β-galactoside (IPTG) and left to grow overnight (22.5°C, 210 rpm) before harvesting by centrifugation (6,239 × g, 13 minutes). Cell pellets containing galectin-3 were resuspended in lysis buffer; 50 mM HEPES (pH 7.8), 300 mM NaCl, cOmplete EDTA-free protease inhibitor cocktail (1 tablet per 100 mL) (Roche), 0.1 mg/mL lysozyme (Sigma-Aldrich) and 10 μg/mL deoxyribonuclease I (DNase1) (Sigma-Aldrich). Cells were lysed with two passes at 28 kpsi using a cell disruptor (Constant Systems), after which the lysate was clarified (53,343 × g, 30 minutes). The supernatant containing soluble galectin-3 was purified on a 5 mL HisTrap HP column (Cytiva) pre-equilibrated with buffer containing 50 mM HEPES (pH 7.8), 300 mM NaCl and 10 mM imidazole. Following binding, the column was washed; 50 mM HEPES (pH 7.8), 300 mM NaCl and 20 mM imidazole. Bound protein was eluted using an imidazole gradient (start concentration: 0 mM, end concentration: 750 mM) with the protein eluting at approximately 300 mM imidazole. Eluted galectin-3 was dialysed overnight against 50 mM HEPES (pH 7.8), 300 mM NaCl, 0.3 mM tris(2-carboxyethyl)phosphine (TCEP). The His-tag was cleaved during dialysis by addition of Human Rhinovirus (HRV) 3C-protease at a 1:100 ratio 3C:galectin-3. Cleaved galectin-3 was purified from non-cleaved materials using reverse immobilised metal affinity chromatography (IMAC) on a pre-equilibrated 5 mL HisTrap HP column (Cytiva). The flow through was collected and the column washed as previous before eluting the bound material; 50 mM HEPES (pH 7.8), 300 mM NaCl and 375 mM imidazole. Samples of cleaved galectin-3 were purified further using size exclusion chromatography (SEC) with a Superdex S75 10/300 column (Cytiva) run at 0.5 mL/min. Fractions of galectin-3 were confirmed using SDS-PAGE analysis confirming presence and purity. The galectin-3 protein purified for SPR studies was highly pure with minimal contaminant.

#### Surface plasmon resonance

All SPR experiments were performed on the Biacore T200 (Cytiva) in running buffer; 0.1 M HEPES (pH 7.4), 1.5 M NaCl and 0.5% surfactant P20 (HBS-P+ Buffer 1 ×) +/-1 mM MnCl_2_. Prior to commencing experiments the instrument was successfully primed and underwent routine normalisation to ensure a uniform signal during the assay. Commercially available recombinant human integrins αvβ1, αvβS and αvβ6 (R&D Systems) or TGF beta receptor II protein (Abcam) were diluted in 10 mM sodium acetate pH 4.0 immobilisation buffer and immobilised onto a Series S Sensor Chip CM5 (Cytiva) via the standard amine coupling protocol. Serial dilutions (2-fold) of galectin-3 in running buffer were injected at a 30 μL/min flow rate, 20°C, contact time: 120 s and dissociation time: 1200 s (integrins) or 300 s (TGFβRII) into flow cells containing approximately 1000 RU of integrin (equating to approximately 135 Rmax) or approximately 400 RU TGFβRII protein (equating to approximately 2S1 Rmax). The sensor chip surface was regenerated with 5 mM ethylenediamine tetraacetic acid (EDTA) for integrin-galectin-3 experiments. All sensorgrams and kinetic plots were baseline-corrected using a flow cell with blank immobilisation prior to data analysis (Figure S-1). Binding data was analysed in GraphPad Prism, if steady-state was reached then the K_d_ value (M) and Bmax were determined by non-linear regression (binding saturation) - one-site specific binding. If steady-state was not reached then the minimum number of binding sites was estimated from the response at the highest analyte concentration tested (5000 nM). This was calculated using the following formula (57):

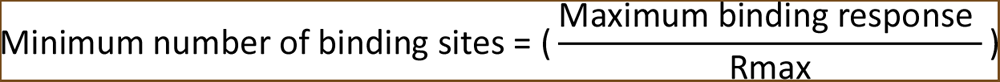

To determine the effects of deglycosylation on galectin-3 binding to integrins or TGFβRII, all glycans were removed from recombinant proteins using protein deglycosylation mix II (New England Biolabs) or N-linked glycans alone removed via PNGase F digestion (New England Biolabs) under non-denaturing conditions. Successful deglycosylation was confirmed by SDS-PAGE and deglycosylated proteins immobilised as described above. The same galectin-3 serial dilution was injected across all flow paths at a 30 μL/min flow rate, 20°C, contact time: 120 s and dissociation time: 600 s (integrins) or 300 s (TGFβRII).

SPR was used to assess the effects of small molecule galectin-3 inhibitors (GB0139 and GB1107) on galectin-3 binding to integrins or TGFβRII. Glycosylated recombinant proteins were immobilised as described above. Galectin-3 inhibitors were serially diluted (2-fold) in running buffer containing 625 nM galectin-3 across the dilution series (stocks of compound in 100% DMSO diluted to a 1.5% final concentration). The serial dilution of compound (+ galectin-3) was injected across all flow paths at a 30 μL/min flow rate, 20°C, contact time: 120 s and dissociation time: 600 s (integrins) or 300 s (TGFβRII). The response values were normalised with respect to the highest RU response. Competition binding curves were analysed in GraphPad Prism and inhibitory concentration 50 (IC50) values determined by non-linear regression analysis (binding saturation) - specific binding with hill slope.

#### Co-immunoprecipitation

Co-IP was performed using the Pierce Crosslink IP Kit (Thermo Scientific). Cells were harvested by scraping in 1X phosphate-buffered saline (PBS), pellets were lysed in ice cold IP lysis/ wash buffer (300 μL/S0 mg wet pellet) and protein concentrations determined by Pierce BCA assay (Thermo Scientific). Lysates (6S0 μg/Co-IP reaction) were pre-cleared using control agarose resin slurry (S2 μL/ Co-IP reaction) for 1 hour at 4°C. The pre-cleared lysates were then incubated at 4°C overnight with protein A/G plus agarose coupled and crosslinked to 10 μg of pulldown antibody: mouse anti-integrin beta 1/CD29 mAb (R&D Systems, Clone No. P5D2), rabbit anti-galectin-3 mAb (Abcam, Clone No. EPR19244), mouse IgG1 isotype control mAb (R&D Systems, Clone No. 11711) or rabbit IgG isotype control mAb (Abcam, Clone No. EPR25A). Flow-throughs and wash steps were collected until confirmation of successful IP. Elution buffer was added to spin columns and eluates analysed by western blot for the presence of antigen. Samples were subjected to SDS-PAGE on a 4-12% Bis-Tris gel (Invitrogen) and transferred onto a PVDF membrane (Invitrogen). After blocking for 2 hours; 5% non-fat milk in PBS plus 0.05% Tween 20 (PBST), the membrane was incubated overnight at 4°C with primary antibody: rabbit anti-galectin-3 mAb (Abcam, Clone No. EPR19244; 1:1000) or mouse anti-integrin beta 1 mAb (Abcam, Clone No. 12G10; 1:1000). After washes (PBST) the membrane was incubated for 1 hour at room temperature with a HRP-conjugated secondary antibody: goat anti-rabbit Immunoglobulins/HRP pAb (Dako; 1:3000) or goat anti-mouse IgG/HRP pAb (Invitrogen; 1:5000). All primary and secondary antibodies used were diluted in blocking buffer. The membrane was then incubated with ECL western blot detection reagent and visualised using chemiluminescent detection.

#### Proximity ligation assay

Cells were growth arrested in serum-free media for 24 hours prior to experiments. After serum starvation, cells were stimulated with 2 ng/mL recombinant human TGF-β1 protein (R&D Systems) for 24 hours and then pre-treated with 0.1-10 μM galectin-3 inhibitor (GB0139) for 20 minutes. PLA was performed using the Duolink In Situ Red Kit Mouse/Rabbit (Sigma-Aldrich). Prior to PLA, cells were fixed with 4% paraformaldehyde (PFA) for 10 minutes and incubated with blocking solution (1X PBS pH 7.4, 10% FBS) for 2 hours at 37°C. Slides were then incubated overnight at 4°C with primary antibody (5 μg/mL): mouse anti-integrin beta 1 mAb (Abcam, Clone No. P5D2), rabbit anti-galectin-3 pAb (Invitrogen), mouse IgG1 isotype control mAb (Dako, Clone No. DAK-GO1) or rabbit IgG isotype control pAb (BD Biosciences, Clone No. Poly1281). After washes (Wash Buffer A), slides were incubated for 2 hours at 37°C with PLA probe solution containing anti-rabbit PLUS and anti-mouse MINUS probes diluted in antibody diluent (1X PBS pH 7.4, 2% FBS, 1% BSA) (1:S). Ligase (1 U/μL) diluted in 1X ligation buffer (1:40) was then added to washed slides (Wash Buffer A) and incubated for 30 minutes at 37°C. Polymerase (10 unit/μL) was diluted in 1X amplification buffer (1:80) and added to slides for 2 hours at 37°C after wash steps (Wash Buffer A). After washes (Wash Buffer B), slides were then stained with DAPI solution and mounted with fluoroshield for imaging (Leica LSM 980 Airyscan 2).

#### Statistical analysis

Data analysed in GraphPad Prism and presented as mean with individual replicates shown.

#### Supporting information

This manuscript contains supporting information and includes additional references (38, 65-67).

## Acknowledgements

This work was supported by the Medical Research Council (MRC IMPACT iCASE Grant: MR/R015813/1) and Galecto Biotech. We would like to thank the patients who consented to provide samples for use in this research. We would also like to thank Professor Jerry Shay (University of Texas, USA) for the gift of iHBECs and Professor Simon Macdonald (University of Nottingham, UK and GSK) for the gift of β1 integrin inhibitor (NOTT199SS). NOTT199SS was identified as part of an MSci Chemistry undergraduate integrin drug discovery collaboration between the School of Chemistry at the University of Nottingham and GSK, supervised by Dr Simon Macdonald (GSK) and Dr Thomas McInally (University of Nottingham). We thank the Advanced Imaging Facility (RRID:SCR_020967) at the University of Leicester for support (BBSRC Grant: BB/S019510/1).

