## Supplemental Table 1 for "Defining the mechanism of galectin-3-mediated TGF-β1 activation and its role in lung fibrosis"

**Table S1. Galectin-1 and Galectin-3 $K_d$ values and cell permeability for Galecto Biotech compounds**

| Compound | Galectin-3 $K_d$ (µM) | CACO-2 (A > B/B > A) Papp (10−6 cm/s) |
|---|---|---|
| GB0139 | 0.0023 | 0.07/0.05 |
| GB1211 | 0.025 | 4.6/30 |
| GB1107 | 0.037 | 15/16 |
| GB0149 | 0.099 | <0.03/<0.07 |
