## Supplemental Figure 1 for "Defining the mechanism of galectin-3-mediated TGF-β1 activation and its role in lung fibrosis"

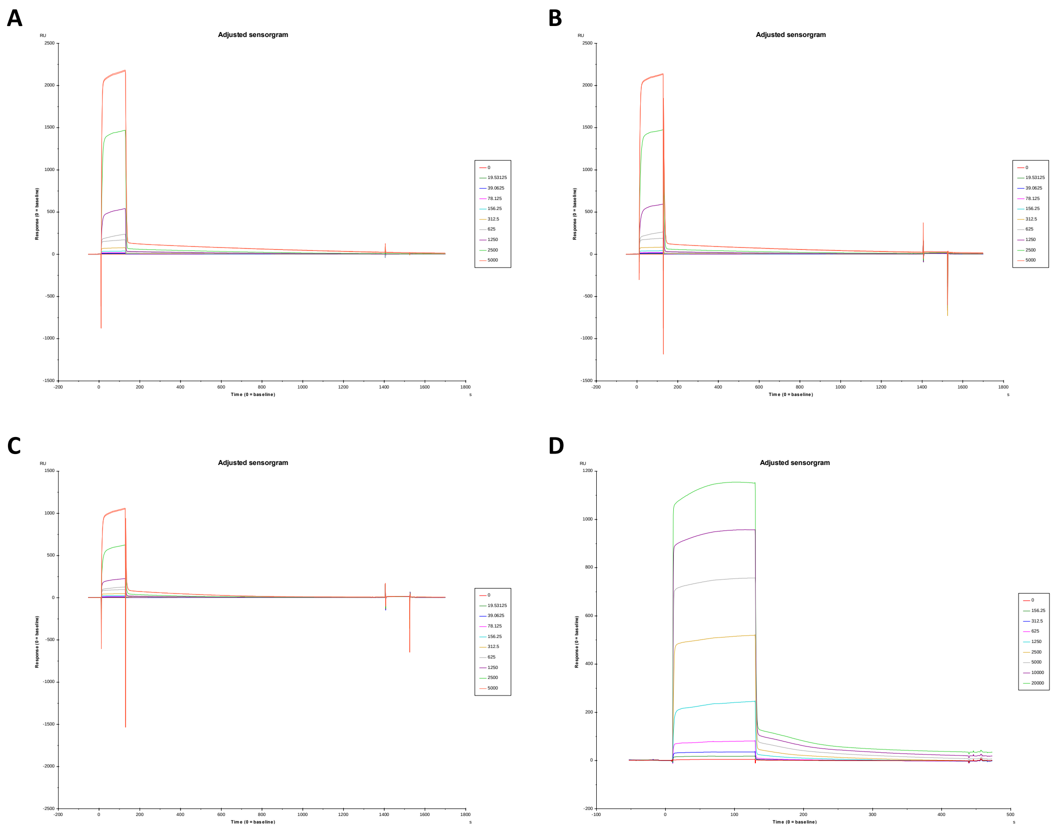


**Figure S1. SPR Sensorgrams for galectin-3 binding to glycosylated αv integrins or TGFβRII**

Baseline-corrected sensorgrams for galectin-3 binding to glycosylated αv integrins: (A) αvβ1, (B) αvβ5 and (C) αvβ6 or (D) the TGFβRII subunit.
